# Community-Based Surveillance for Highly Pathogenic Avian Influenza Viruses among Deceased Birds

**DOI:** 10.64898/2026.03.06.710164

**Authors:** Lyudmyla V. Marushchak, Caitlin J. Cotter, Judith U. Oguzie, Philip K. Keiser, Thang Nguyen-Tien, Jessica Rodriguez, Ismaila Shittu, Claudia M. Trujillo-Vargas, Amanda E. Wolff, Shivonne M. Ryans, Robert D. Kaufman, Jillian D. Clack, Susan L. F. McLellan, Gene G. Olinger, Gregory C. Gray

**Affiliations:** Division of Infectious Diseases, Department of Medicine, University of Texas Medical Branch, Galveston, TX, USA; Department of Epidemiology, School of Public Health and Health Professions, University of Texas Medical Branch, Galveston, Texas, USA; Institute for Human Infections and Immunity, University of Texas Medical Branch, Galveston, TX, USA; Department of Microbiology and Immunology, University of Texas Medical Branch, Galveston, TX, USA; Galveston County Health District, Galveston, Texas, USA; Galveston County Health District Animal Services team, Galveston, Texas, USA; Special Pathogens Excellence in Clinical Treatment, Readiness, and Education, Galveston, Texas, USA; Galveston National Laboratory, University of Texas Medical Branch, Galveston, Texas, USA

**Keywords:** avian influenza, Texas, wild birds, surveillance, epidemiology

## Abstract

Highly pathogenic avian influenza (HPAI) viruses of H5N1 clade 2.3.4.4b, are spreading worldwide, posing a threat to wildlife, domestic animals, and humans. In 2025, a multidisciplinary collaboration for HPAI H5N1 surveillance among birds within Galveston County, Texas, was initiated. Between November and December 2025, oropharyngeal and cloacal swabs were collected from wild and domestic birds reported as dead or dying by Galveston County residents. Specimens were studied with molecular assays, Sanger sequencing, virus isolation, and next-generation sequencing. Molecular evidence of HPAI H5N1 was detected in 7 of 10 (70%) birds, and the virus was successfully cultured in MDCK cells. Next-generation sequencing analysis of eight influenza A genome segments demonstrated a 4:4 gene segment reassortant constellation within clade 2.3.4.4b, consistent with genotype D1.1. Community members exposed to HPAI were offered antiviral prophylaxis. No human infections were identified. This surveillance demonstrates that community involvement combined with cross-sectoral collaboration can ensure rapid detection and characterization of circulating avian influenza viruses. Sustained local surveillance is essential for early warning, risk assessment, and prevention of virus spread to poultry, mammals, and humans.

## Introduction

Highly pathogenic avian influenza (HPAI) viruses are widely disseminated via migratory wild birds. Since 2020, the spread of various clades of HPAI H5 viruses has escalated into a global panzootic [1]. In 2021, the Eurasian strain of H5N1 (clade 2.3.4.4b) was detected among wild birds in North America [2; 3]. Large outbreaks in U.S. poultry populations occurred in 2022, with migratory birds playing a critical role in long-distance virus dissemination. Strains of HPAI H5N1 additionally have spread from U.S. wild birds to an increasing range of mammalian species worldwide [4; 5]. In 2024, HPAI H5N1 clade 2.3.4.4b spread to U.S. dairy cows with sporadic human infections documented among U.S. dairy and poultry workers [6; 7]. As HPAI H5N1 strains continue to infect a range of mammalian species, including causing life-threatening disease in domestic cats [8], the increasing frequency and geographic spread of these epizootics have heightened concerns regarding the virus’ potential for further viral adaptations in strains transmitted among both avian and mammalian host to pose a greater threat to both humans and livestock.

Widespread transmission events in avian populations, at times associated with significant population die-off events caused by H5N1 threaten ecosystem stability, biodiversity conservation, and species recovery, and continued transmission to livestock underscores potential risk to human communities. Mitigating these threats requires coordinated wildlife surveillance, rapid genomic characterization of circulating viruses to evaluate viral mutations enabling species spillover, and collaboration across animal, environmental, and public health sectors within a One Health framework. Continued surveillance for viral adaptation in circulating strains of HPAI H5N1, particularly along migratory flyways that facilitate viral movement across regions is paramount.

Galveston County, Texas, USA, is a stopover point for migratory birds along the Central migratory Flyway, providing essential habitats for species traveling between breeding grounds in North America and wintering areas in Central and South America [9]. Its diverse coastal habitats support more than 300 bird species annually. The ecological importance of Galveston County for migratory birds increases both the likelihood of virus introduction and the need for vigilant avian influenza surveillance in wild bird populations in the area.

Given these concerns and limitations of public surveillance systems for wildlife disease, a HPAI H5N1 surveillance program was initiated through a multidisciplinary collaboration between Galveston County Health District (GCHD), the University of Texas Medical Branch (UTMB) Schools of Medicine and Public and Population Health, Galveston County Health District Animal Services team, UTMB’s Special Pathogens Excellence in Clinical Treatment, Readiness, and Education Program (SPECTRE), the Galveston National Laboratory (GNL), and the general public. This collaboration sought to strengthen local capacity for the rapid detection and characterization of avian influenza viruses in dead wild or domestic birds encountered by the general public in Galveston County. This report summarizes the results from the first collection of deceased bird specimens sampled through the program.

## Methods

### Planning the surveillance

In February 2025, multidisciplinary partners were invited to meet with GCHD to develop a citizen science-based HPAI H5N1 surveillance program and to seek UTMB institutional animal care and use committee (IACUC) review of program protocols. In September 2025, UTMB IACUC determined that activities involving deceased carcasses did not require formal ethical review, and surveillance plans were set in motion. UTMB partners met with the Galveston County Health District Animal Services team for training on public communication, personal protective equipment (PPE) use, specimen and data collection, field specimen packaging, specimen delivery to UTMB’s One Health Research and Training Laboratory (OHRTL), and communication of results. UTMB and GCHD provided PPE, materials for specimen collection, cold packs, and transport containers to the GCHD Animal Resources team. GCHD communication professionals, in collaboration with UTMB faculty and students, developed flyers for the general public, directing citizens to contact GCHD to report findings of dead wild birds or sightings of birds with neurological symptoms.

### Surveillance Strategy

When a report of a dead animal was received by GCHD, specimens were collected by GCHD’s Animal Services team and delivered to UTMB’s OHRTL. Samples were studied for molecular evidence of influenza A virus (IAV). Positive samples were further characterized with a battery of additional laboratory studies, which often included additional RT-qPCR for H5 and H7, virus isolation in cell culture, Sanger sequencing, and next-generation sequencing (**Figure 1**). Laboratory reports were rapidly communicated to the GCHD Authority, Dr. Phil Keiser, and other members of the surveillance team. Subsequently, the GCHD issued recommendations to the community regarding influenza prophylaxis for individuals with potential exposure as well as monitoring recommendations for exposed household animals.

**Fig. 1.**
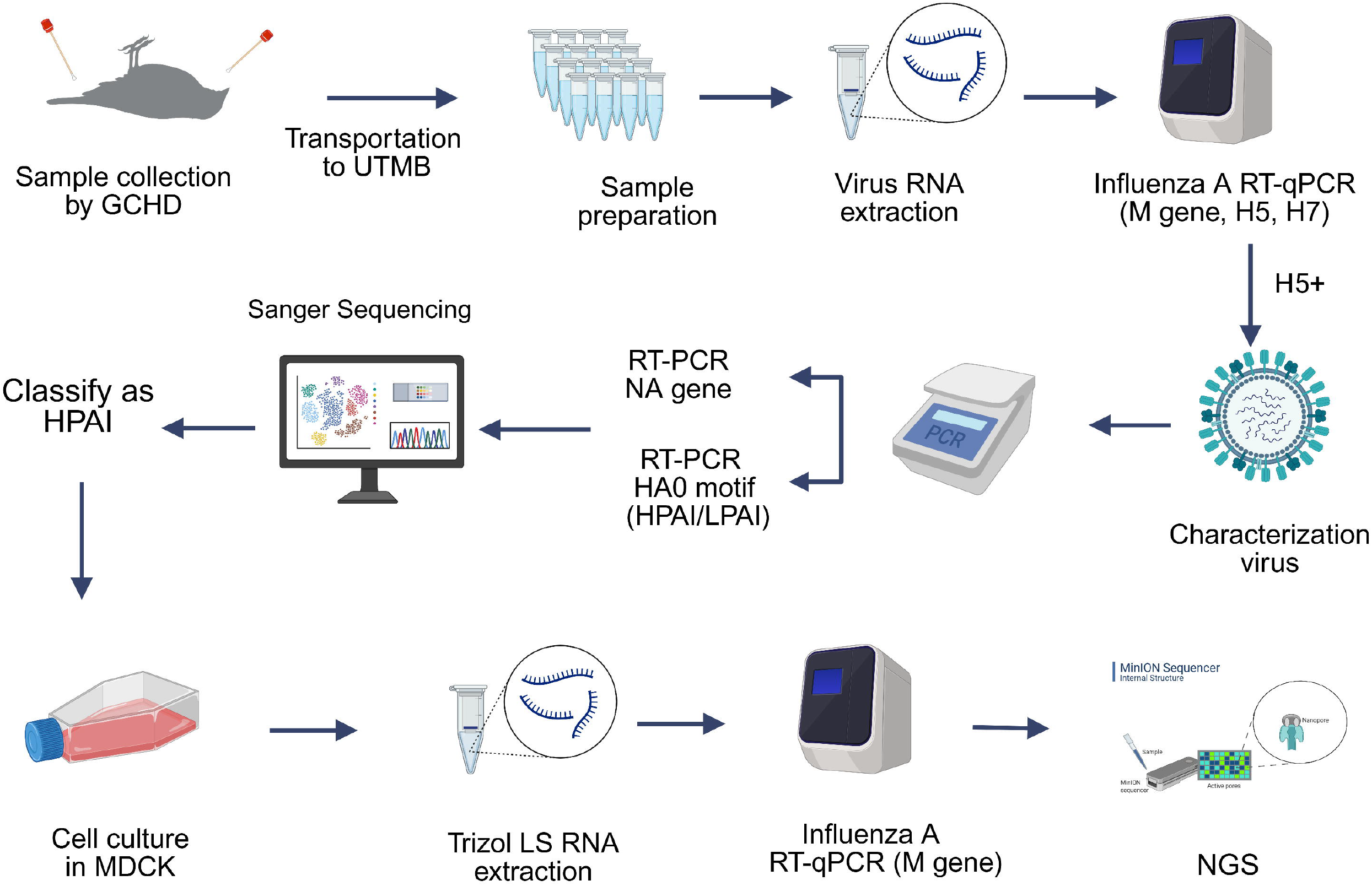
The workflow implemented to identify and characterize influenza A virus in samples collected from deceased birds. The testing methods include RNA extraction, RT-qPCR and RT-PCR, Sanger sequencing, and next-generation sequencing. This figure was created with BioRender.com. GCHD, Galveston County Health District; UTMB, University of Texas Medical Branch; RNA, ribonucleic acid; HPAI, highly pathogenic avian influenza; LPAI, low pathogenic avian influenza; RT-qPCR, quantitative RT-PCR; RT-PCR, reverse transcription PCR; M gene, Influenza A Matrix gene; H5, Influenza A hemagglutinin subtype 5; H7, Influenza A hemagglutinin subtype 7; NA, Influenza A neuraminidase gene; HA0, hemagglutinin (HA0) cleavage site of influenza A; MDCK, Madin-Darby canine kidney cells; NGS, Next Generation Sequencing.

### Sample Collection

The GCHD Animal Resources team collected oropharyngeal and cloacal swab specimens from dead wild birds and completed a field collection form. Each swab specimen was placed in 2 mL of sterile viral transport medium (VTM; Rocky Mountain Biological, Crested Butte, CO) and sent within 24 hours to the OHRTL, UTMB at Galveston, Texas, USA while maintaining a cold chain between 2 °C and 8 °C. At UTMB, the samples were aliquoted and stored at −80 °C until tested.

### Molecular Detection

At UTMB, virus inoculation and virus inactivation were performed in a BSL3 enhanced (BSL3E) laboratory following BSL3E work practices. The TRIzol™ LS Reagent (Invitrogen, Waltham, MA) was used for virus inactivation. Samples were then transferred to a BSL2 laboratory for RNA extraction and characterization. Viral RNA was extracted from the swab specimens using QIAamp Viral RNA Mini Kit (Qiagen, Hilden, Germany) following manufacturer’s instructions using the QIAcube Connect RNA/DNA Extraction Instrument (Qiagen).

All specimens were screened for influenza A virus (Matrix gene) using reverse transcription quantitative real-time polymerase chain reaction (RT-qPCR) and positive samples were subsequently subtyped for H5 and H7 using protocols recommended by the World Health Organization [10; 11]. Samples positive for H5 were further characterized to distinguish highly pathogenic avian influenza (HPAI) from low pathogenic avian influenza (LPAI) virus subtypes using RT-PCR to target the hemagglutinin (HA0) cleavage site for amplicon sequencing [12]. Additionally, influenza A/H5 positive samples were tested for influenza A neuraminidase (NA) genes by conventional RT-PCR [13]. The amplicons were analyzed by electrophoresis on a 2% agarose gel. Amplicons with the expected targeted molecular weights for the NA gene and HA0 cleavage site were sent to GENEWIZ from Azenta Life Sciences (https://www.genewiz.com) for Sanger sequencing.

Influenza A virus positive samples were subsequently transferred from BSL2 to the BSL3E laboratory for long-term storage.

### Sanger sequencing

Amplicons of the expected size for the HA0 cleavage site and NA gene were further studied with Sanger sequencing by GENEWIZ. The Basic Local Alignment Search Tool (BLAST) available on the National Center for Biotechnology Information (NCBI) platform was used for rapid sequence comparison and the determination of species identity based on the highest sequence similarity. The resulting sequence data were studied using Geneious Prime^®^ version 2026.0.2 for sequence editing and alignment.

The EMBOSS SIXPACK (https://www.ebi.ac.uk) was used to generate six-frame translations of the nucleotide sequences to differentiate HPAI from LPAI viruses. For downstream analysis, translations containing the minimum number of open reading frames (ORFs) were selected.

### Cell culture: Virus Isolation

Madin-Darby canine kidney cells (MDCK; ATCC, cat no. CRL-CCL34) were used for influenza A culture efforts. The MDCK cells were grown in Dulbecco’s Modified Eagle Medium (DMEM) containing high glucose and L glutamine (ThermoFisher Scientific cat no. 11965118) supplemented with 10 % fetal bovine serum (FBS, ThermoFisher Scientific cat no. 26140–079) and 2 % Antibiotic-Antimycotic (100X) (ThermoFisher Scientific, cat no. 15240–062).The cells were cultured as monolayers in 6-well plates at 37 °C with 5 % CO2.

Original oropharyngeal and cloacal swab specimens were filtered using 0.45 μm pore-size filters (Millipore Sigma, Ireland) and then 0.2 ml aliquots were inoculated onto the cells with an addition of 0.8 mL of serum-free maintenance medium (DMEM, 2 % Antibiotic-Antimycotic). Cells were observed microscopically daily for cytopathic effect (CPE) for 3-4 days.

### Whole Genome Amplification and Nanopore Sequencing

Influenza A positive isolates were characterized by next-generation sequencing (NGS) of whole genome segments using the Oxford Nanopore MinION platform [14]. The resulting mRT-PCR products were prepared for sequencing with the Native Barcoding Kit 96 V14 (SQK-NBD114.96, Oxford Nanopore Technologies (ONT), England). Sequencing was performed on R10.4.1 MinION flow cells (FLO-MIN114, ONT, England) on the MinION Mk1D platform.

Raw sequencing data were basecalled and demultiplexed using the high accuracy basecalling mode in the ONT MinKNOW software. Sequence reads were analyzed on the CZ ID platform (czid.org) to identify IAV contigs. Viral subtypes were confirmed using the Genome Detective Influenza Typing Tool (https://www.genomedetective.com/app/typingtool/iav/) and NCBI BLAST against the Influenza virus database (https://www.ncbi.nlm.nih.gov/genomes/FLU/Database/nph-select.cgi). Consensus genomes were generated via Minimap 2.24 and mutation analysis was carried out on the FluSurver (http://flusurver.bii.a-star.edu.sg/). We performed phylogenetic comparisons of related viruses in GenBank and Global Initiative on Sharing All Influenza Data (GISAID EpiFlu™) (https://gisaid.org) [15]. Sequences were aligned using MAFFT v7.490 [16; 17]. Maximum likelihood phylogenetic trees were constructed using IQ-TREE [18] with 1,000 bootstrap replicates, and the best-fit substitution model was automatically selected by ModelFinder. The resulting trees were visualized and annotated using FigTree v1.4.4 (http://tree.bio.ed.ac.uk/software/figtree/). The H5N1 sequences were assigned clades using the Nextclade (https://clades.nextstrain.org/). GenoFLU v1.06 was used for genotyping.

## Results

During November and December 2025, oropharyngeal and cloacal swabs from 10 dead birds (**Table 1**). Overall, specimens were collected from four Muscovy ducks (*Cairina moschata*) in Dickinson, one chicken (*Gallus gallus domesticus*) in Santa Fe, one duck hybrid in Santa Fe, one Woodcock-like bird (*Scolopax minor*) in La Marque, one Cattle Egret *(Bubulcus ibis)* in Texas City, and one wild bird in La Marque. Residents often reported evidence of neurological signs of illness in the birds before they died.

**Table 1.** Summary of the influenza A virus detections and molecular assay results from deceased bird swabs collected in Galveston County, Texas during November to December 2025.

| Bird Type | Sample ID | Sample Type | City | Influenza A (M gene) RT-qPCR | Influenza A (H5) RT-qPCR | HA0 motif RT-PCR HPAI<sup>a</sup>/LPAI<sup>b</sup> | Influenza A (NA<sup>c</sup> gene) RT-PCR |
| --- | --- | --- | --- | --- | --- | --- | --- |
|  |  |  |  | Ct value | Ct value | Sanger result | Sanger result |
| Wild bird | GCAI004_1 | oropharyngeal | La Marque | Negative | Negative | TNP<sup>d</sup> | TNP<sup>d</sup> |
|  | GCAI004_2 | cloacal |  | Negative | Negative | TNP<sup>d</sup> | TNP<sup>d</sup> |
| Muscovy duck | GCAI005_1 | oropharyngeal | Dickinson | 21.09 | 24.01 | HPAI | N1 |
|  | GCAI005_2 | cloacal |  | 29.67 | 23.44 | HPAI | N1 |
| Muscovy duck | GCAI006_1 | oropharyngeal | Dickinson | 22.92 | 24.46 | HPAI | N1 |
|  | GCAI006_2 | cloacal |  | 23.08 | 24.49 | HPAI | N1 |
| Muscovy duck | GCAI020_1 | oropharyngeal | Dickinson | 18.97 | 19.98 | HPAI | N1 |
|  | GCAI020_2 | cloacal |  | 23.35 | 25.28 | HPAI | N1 |
| Muscovy duck | GCAI021_1 | oropharyngeal | Dickinson | 22.05 | 23.21 | HPAI | N1 |
|  | GCAI021_2 | cloacal |  | 22.21 | 24.31 | HPAI | N1 |
| Chicken | GCAI014_1 | oropharyngeal | Santa Fe | 22.95 | 25.85 | HPAI | N1 |
|  | GCAI014_2 | cloacal |  | 19.80 | 23.66 | HPAI | N1 |
| Duck hybrid | GCAI026_1 | oropharyngeal | Santa Fe | 30.20 | 32.56 | HPAI | N1 |
|  | GCAI026_2 | cloacal |  | 16.17 | 19.20 | HPAI | N1 |
| Cattle Egret | GCAI012_1 | oropharyngeal | Texas City | Negative | Negative | TNP<sup>d</sup> | TNP<sup>d</sup> |
|  | GCAI012_2 | cloacal |  | Negative | Negative | TNP<sup>d</sup> | TNP<sup>d</sup> |
| Woodcock | GCAI011_1 | oropharyngeal | La Marque | Negative | Negative | TNP<sup>d</sup> | TNP<sup>d</sup> |
|  | GCAI011_2 | cloacal |  | Negative | Negative | TNP<sup>d</sup> | TNP<sup>d</sup> |
| Yellow Caracara | GCAI018_1 | oropharyngeal | Texas City | 25.12 | 29.23 | HPAI | N1 |
|  | GCAI018_2 | cloacal |  | 31.06 | 35.53 | HPAI | N1 |

<sup>a</sup> Highly pathogenic avian influenza, <sup>b</sup> Low pathogenic avian influenza, <sup>c</sup> Neuraminidase gene, <sup>d</sup> test not performed

Molecular evidence of influenza A virus was detected in 7 of the 10 (70%) birds. The influenza positive samples were confirmed as subtype H5 by RT-qPCR. The multibasic HA cleavage site, consistent with HPAI, was detected by Sanger sequencing in all. Amplicon sequencing for the NA gene demonstrated that all HPAI H5 positive specimens had evidence of N1. All samples were negative for influenza A subtype H7. In UTMB’s BSL3E laboratory, the HPAI H5N1 virus was successfully cultured in MDCK cells from all seven influenza A positive birds.

Analysis of all eight genome segments revealed a 4:4 reassortant constellation within clade 2.3.4.4b, consistent with genotype D1.1. Additionally, this genotype was confirmed by HA phylogenetic analysis (**Figure 2**). Specifically, D1.1 has been shown to have 4 of the 8 genes, (PB1, HA, MP, and NS) of the Eurasian avian lineage, and the other 4 gene (PB2, PA, NP, and NA (N1)) of North American low-pathogenicity avian influenza (LPAIV) lineages, matching the expected D1.1 reassortment pattern [19]. Mutation analysis of the eight genome segments identified notable mutations in segments 1 (PB2), 2 (PB1), 6 (NA), and 8 (NS1) (**Table 2**). These multiple amino acid substitutions have been previously associated with virulence, host adaptation, antigenic evolution, and antiviral susceptibility. All genome sequences generated in this study have been deposited in GenBank under accession numbers PX995912–PX996007 and GISAID EpiFlu™ under accession numbers EPI_ISL_5152143–EPI_ISL_5152238.

**Fig. 2.**
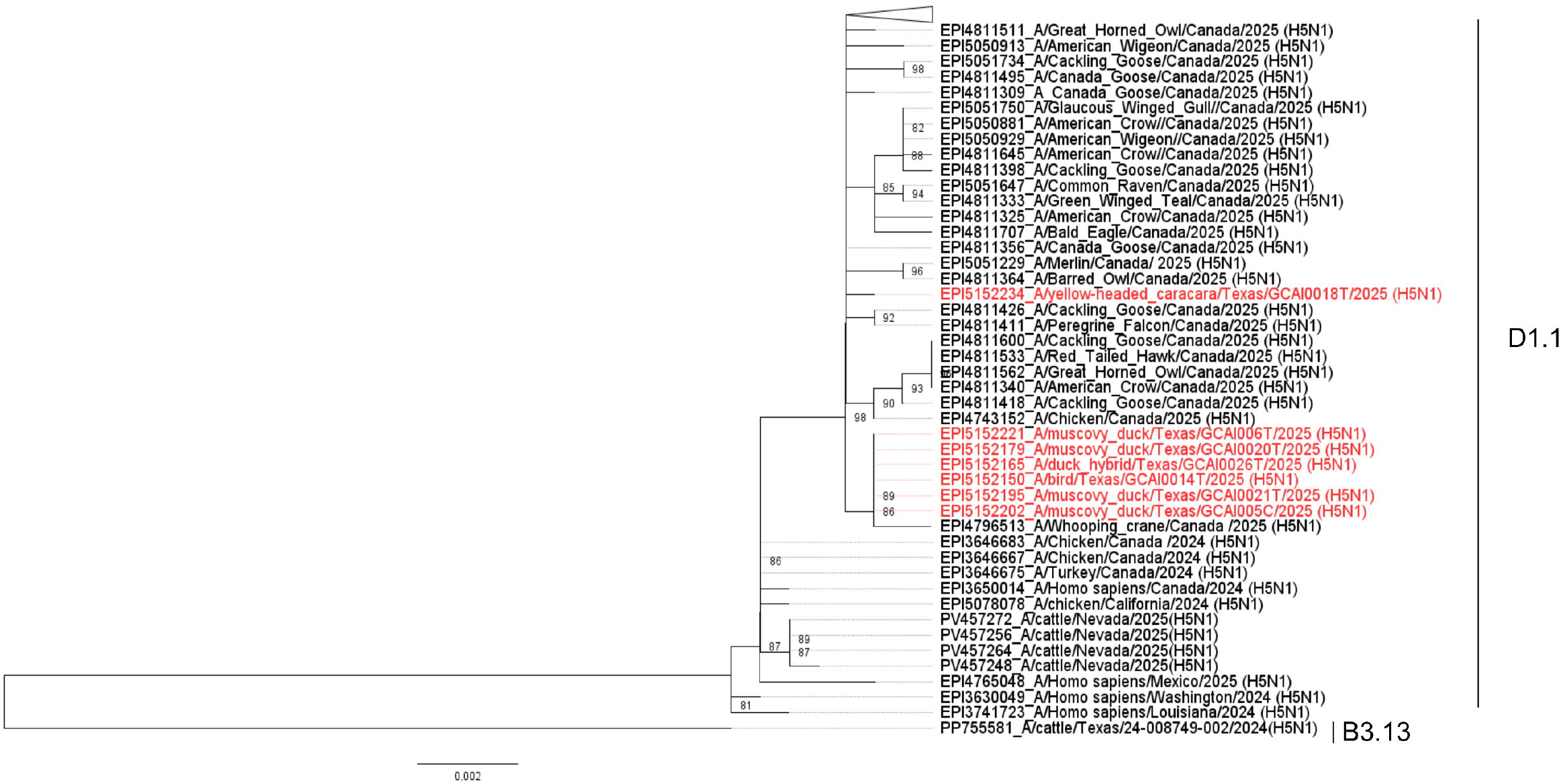
Phylogenetic tree of hemagglutinin (HA) gene sequences of H5N1 influenza A viruses detected in this study. Maximum-likelihood tree inferred from sequences generated in this study (highlighted in red). Closely related reference sequences were downloaded from National Center for Biotechnology Information (NCBI) and Global Initiative on Sharing All Influenza Data (GISAID).

**Table. 2.**
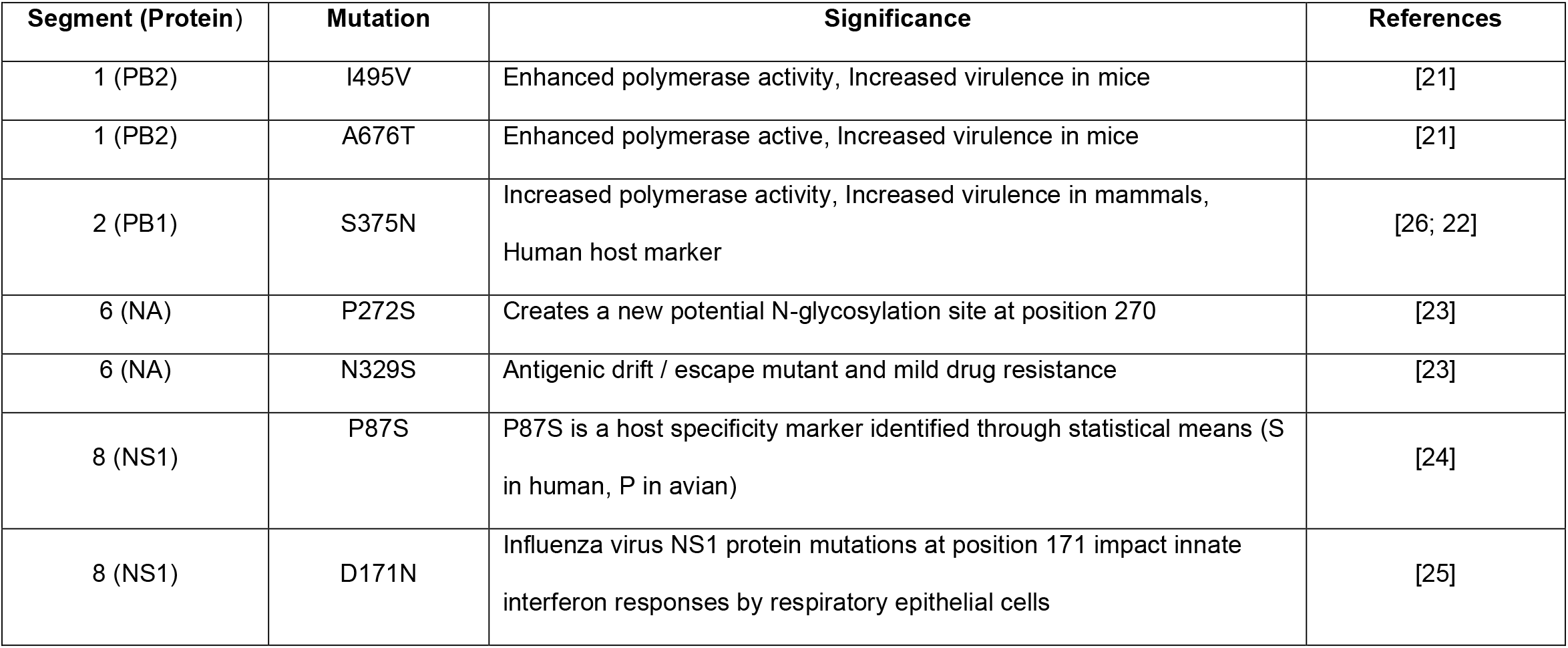
Summary of relevant mutations in the genome of highly pathogenic avian influenzas A viruses of H5N1 clade 2.3.4.4b isolated from dead birds in Galveston County Texas, November through December 2025.

| Segment (Protein) | Mutation | Significance | References |
| --- | --- | --- | --- |
| 1 (PB2) | I495V | Enhanced polymerase activity, Increased virulence in mice | [21] |
| 1 (PB2) | A676T | Enhanced polymerase active, Increased virulence in mice | [21] |
| 2 (PB1) | S375N | Increased polymerase activity, Increased virulence in mammals, Human host marker | [26; 22] |
| 6 (NA) | P272S | Creates a new potential N-glycosylation site at position 270 | [23] |
| 6 (NA) | N329S | Antigenic drift / escape mutant and mild drug resistance | [23] |
| 8 (NS1) | P87S | P87S is a host specificity marker identified through statistical means (S in human, P in avian) | [24] |
| 8 (NS1) | D171N | Influenza virus NS1 protein mutations at position 171 impact innate interferon responses by respiratory epithelial cells | [25] |

Ten community members reported direct contact with the infected birds. Among these exposed individuals, eight opted to receive oseltamivir (Tamiflu®) post-exposure prophylaxis. Exposed individuals who subsequently developed respiratory symptoms had nasal swabs collected for rapid influenza virus testing; all were found to be negative.

## Discussion

This project demonstrates the feasibility of local multisector surveillance collaborations. This collaboration highlights the need to develop alternate strategies for emerging pathogens surveillance, as called for by a recent commentary by RB Shpiner [20].

In this surveillance work, we detected avian influenza A H5N1 viruses consistent with genotype D1.1 genome constellation from dead birds. Identification of a D1.1-like genome constellation in the context of avian mortality is epidemiologically important, as it indicates ongoing circulation of this reassortant virus in wild bird populations and underscores the role of wild birds as both maintenance hosts and sources of repeated introductions into new settings.

Beyond identification of the 4:4 reassortant genotype D1.1 constellation, the detection of specific molecular markers associated with mammalian adaptation and immune modulation is epidemiologically significant. Substitutions in the polymerase genes (PB2 I495V, A676T; PB1 S375N) have been experimentally linked to enhanced polymerase activity and increased virulence in mammalian hosts, suggesting potential for improved replication efficiency outside avian species [21; 22]. NA substitutions (P272S and N329S) may influence antigenicity and antiviral susceptibility [23], while NS1 D171N and NS1 P87S may contribute to modulation of innate immune responses and immune evasion [24; 25]. Although no zoonotic transmission was detected in this investigation, the presence of these mutations within wild bird–derived H5N1 viruses underscores the dynamic evolutionary landscape of clade 2.3.4.4b viruses. Continuous genomic monitoring is therefore essential to detect accumulation of adaptive mutations that could increase pathogenicity, transmissibility, or zoonotic risk.

From a One Health perspective, confirmation of D1.1 in dead birds also serves as an early warning for nearby poultry operations and the potential of mammalian spillover, given that carcasses and contaminated environments can facilitate the exposure of scavengers, predators, and peridomestic animals. Accordingly, surveillance is essential to monitor the emergence and spread of reassortant genotypes and to strengthen ongoing risk assessment efforts.

## Acknowledgements

We thank the following participants in the “*Community-Based Partnership for Highly Pathogenic Avian Influenza Viruses among Deceased Wild Birds, Galveston County, Texas” include:* Galveston County Health District: Philip K. Keiser, Amanda E. Wolff; Galveston County Animal Services: Shivonne M. Ryans, Robert D. Kaufman, Alexandra Sierra, Craig Mixson, Dontrae Smith, Emily Barnes, Gretchen Gray; Galveston National Laboratory: Gene G. Olinger; Division of Infectious Diseases, Department of Medicine, School of Medicine UTMB: Lyudmyla V. Marushchak, Judith U. Oguzie, Thang Nguyen-Tien, Jessica Rodriguez, Ismaila Shittu, Claudia M. Trujillo-Vargas, Susan L. F. McLellan, and Corri B. Levine; Department of Epidemiology, School of Public and Population Health UTMB: Caitlin J. Cotter, Jillian Clack; UTMB’s Special Pathogens Excellence in Clinical Treatment: Susan L. F. McLellan, Corri B. Levine. We sincerely thank all data contributors, including the authors and originating laboratories that collected the specimens and the submitting laboratories that generated the genetic sequences and metadata and made them available through the GISAID Initiative (https://gisaid.org/).

## Author contributions

Conceptualization: P.K.K., C.J.C., G.C.G.; Laboratory methodology: L.V.M., J.U.O.; Investigation: L.V.M., J.U.O., T.N.T., J.R., I.S., G.C.G.; Data analysis: L.V.M., J.U.O.; Visualization: L.V.M., J.U.O.; Original draft.: L.V.M., G.C.G., J.U.O., J.R.; Revision: L.V.M., C.J.C.,J.U.O., P.K.K., T.N.T., J.R., I.S., C.T.V., A.E.W., S.M.R., R.D.K., J.D.C., S.L.F.M., G.G.O., G.C.G.; Project administration: G.C.G; Funding acquisition: G.C.G.; Supervision: G.C.G. All authors read and approved the final version of the manuscript.

## Data availability

Data needed to evaluate the conclusions in the manuscript are present in the manuscript. Additional data or specimens may be requested from the corresponding author. The sharing of additional data or specimens will require the signing of a materials transfer agreement.

## Ethical Statement

This surveillance was reviewed by UTMB’s Chair of the Institutional Animal Care and Use Committee. Full committee review was not warranted as: *“No live vertebrate animals will be enrolled, handled, restrained, euthanized, or otherwise used. All specimens will be obtained only from animals found deceased and therefore fall outside the scope of federal and institutional regulations requiring IACUC oversight*.*”*

